# Comparison of shear stress patterns by the established and advanced reconstruction method incorporating side branches to predict plaque progression

**DOI:** 10.1101/2023.04.05.535789

**Authors:** Anantharaman Ramasamy, Lorenz Räber, Ibrahim Halil Tanboga, Hannah Safi, Michalis Hadjiandreou, Antonis Sakellarios, Pieter Kitslaar, Jouke Dijkstra, Flavio G Biccirè, Tom Crake, Lampros K Michalis, Dimitrios Fotiadis, Stephan Windecker, Yao-Jun Zhang, Anthony Mathur, Andreas Baumbach, Ryo Torii, Christos V Bourantas

**Author notes:** Address for correspondence Christos Bourantas MD PhD Consultant Cardiologist, Barts Heart Centre, Barts Heart Centre, West Smithfield, London EC1A 7BE. Funding: AR, AM, AB and CVB are funded by Barts NIHR Biomedical Research Centre, London, UK. Conflicts of interests: All authors have no conflicts of interests to declare.

## Abstract

**Background:** Complete vessel reconstruction (CVR) with incorporation of side branches is essential for accurate evaluation of wall shear stress (WSS) distribution. However, CVR is time consuming and blood flow simulation is computationally expensive, while there is no evidence that WSS computed by CVR, enables better prediction of disease progression compared to WSS derived from the conventional single vessel reconstruction (SVR). We aim to compare the WSS in models reconstructed using the CVR and SVR methods and examine its ability to predict disease progression.

**Methods:** Patients who had baseline and 13-months follow-up intravascular ultrasound (IVUS) imaging (n=19 vessels), and with neoatherosclerotic lesions (n=13 vessels) on optical coherence tomography (OCT) were included in the present analysis. All the studied vessels had at least one side branch with diameter >1mm. 3-dimensional (3D) CVR and SVR were performed and time averaged (TAWSS) and multidirectional WSS were computed using pulsatile blood flow simulation and the performance of both methods in predicting disease progression in IVUS and OCT models were assessed.

**Results:** The incorporation of side branches in 3D geometry resulted in lower TAWSS in the IVUS (0.821 vs 1.698Pa, p<0.001) and OCT-based reconstructions (0.682 vs 1.325Pa, p<0.001) and influenced the multidirectional WSS distribution. In native segments, WSS metrics estimated by the CVR enabled better prediction of the lumen and plaque area and burden at follow-up than SVR and disease progression defined as decrease in lumen area and increase in plaque burden (AUC CVR 0.712 vs SVR 0.554). In stented segments, multidirectional WSS was associated with neointima area in both CVR and SVR methods, but TAWSS was only a predictor of neointima area in the CVR method.

**Conclusions:** The incorporation of side branches in vessel reconstruction influences WSS distribution and enables more accurate prediction of disease progression in native and stented segments than SVR modelling.

**Highlights:**

- Complete vessel reconstruction (CVR) with incorporation of vessel side branches has been proposed for accurate evaluation of wall shear stress (WSS) distribution compared to the traditional single vessel reconstruction (SVR) method; however, there are no studies comparing the performance of the WSS metrics derived by these methods in predicting atherosclerotic evolution.
- In vessels with large side branches, the incorporation of the side branches in the vessel geometry reconstructed from angiographic and intravascular imaging data resulted in lower time averaged wall shear stress (TAWSS) and influenced the multidirectional WSS estimations compared to the models reconstructed without the side branches.
- The WSS metrics estimated in the CVR models enabled better prediction of atherosclerotic disease progression at 13-months follow-up on IVUS than the WSS derived by the SVR.
- In stented vessels, all the WSS metrics in the CVR and the multidirectional WSS in SVR were associated with neointima tissue development; however, both approaches showed limited efficacy in predicting neointima proliferation.

## Introduction

Experimental and clinical studies suggest that local haemodynamic forces distribution is an instigator of vascular inflammation and vulnerable plaque formation and plays a pivotal role on plaque destabilisation and rupture.^1–4^ Traditionally, in clinical studies vessel three-dimensional (3D) reconstruction is performed by fusing high resolution intravascular imaging and angiographic data. However, all the methodologies developed for this purpose make assumptions in the co-registration of the two modalities and the estimation of the orientation of the intravascular images onto the vessel centreline and most of them do not incorporate side branches in the 3D model that can affect flow patterns.^5^ Despite these assumptions, the fusion of angiography and intravascular imaging is regarded as the reference standard for assessing flow patterns and is superior to quantitative coronary angiography (QCA) or coronary computed tomography angiography, as invasive imaging allows detailed assessment of lumen geometry and accurate computation of plaque burden and characterisation of the plaque composition.^6–8^

The methodology used to co-register intravascular imaging and angiographic data on wall shear stress (WSS) values has been recently tested in a study comparing WSS distribution in models that included and those that did not include vessels’ side branches.^9^ The authors reported significant differences in the WSS values in the models generated by the two reconstruction methods. This was attributed to the flowing blood being diverted to side branches resulting in a reduction of the flow in the main vessel and thus, in lower WSS in the geometries with side branches. Therefore, a recently published consensus document on the assessment of WSS in coronary artery reconstructions advocates the incorporation of the side branches in 3D vessel geometry.^10^ However, the methodology developed for complete vessel modelling from intravascular imaging and X-ray data makes approximations in the co-registration of the two datasets, is time consuming and the computational fluid dynamic (CFD) analysis of the generated geometries is computationally expensive. Today, it is unclear whether the CFD analysis in models incorporating side branches enables better prediction of atherosclerotic evolution than the CFD in models which only include the main vessel.^11^

The aim of this study is to compare the time averaged (TA) WSS and multidirectional WSS distribution in models with and without vessels’ side branches and examine whether the local hemodynamic forces distribution in complex geometries including side branches enables better prediction of disease progression in native and stented segments.

## Methods

### Study design

We analysed data from the IBIS-4 study, a multi-modality intravascular imaging study that examined the effects of aggressive statin therapy on plaque evolution in patients presenting with ST-elevation myocardial infarction (NCT00962416). The detailed study design, the inclusion and exclusion criteria, and study endpoints have been reported elsewhere.^12^ In the present sub-study, we included only patients that had biplane coronary angiography at the time of intravascular imaging, and at least a segment with a side branch with diameter >1mm that was assessed by intravascular ultrasound virtual histology imaging (IVUS-VH) at baseline and 13-months follow-up. Short segments <15mm and cases with IVUS hang-up during pullback and those with extensive foreshortening or overlapping on angiography were excluded.

In addition, we analysed angiographic and OCT data acquired from patients that presented with acute coronary syndrome attributed to a neoatherosclerotic lesions that were recruited in a recent study that examined the implication of the ESS distribution on neoatherosclerotic lesion formation and destabilization. The study details and analysis protocol has been described previously.^13^ In the present study we included, only patients that had a stented segments with at least one side branch with a diameter >1mm.

### Data acquisition and image analysis

#### IVUS imaging

IVUS imaging was performed with a 20MHz IVUS catheter (Eagle Eye, Volcano Cooperation, Rancho Cordova, CA) using a motorised pullback speed of 0.5mm/s following administration of intracoronary nitroglycerine. Images were acquired at 30 frames per second (fps) and saved for offline analysis. In the IVUS-VH images acquired at baseline and follow-up, the most proximal and distal side branch that was visible in both examinations and in coronary angiography were identified and used to define the segment of interest. The lumen and external elastic membrane (EEM) were annotated in every frame (approximately 0.4mm) of the segment of interest. Volcano’s proprietary VH technology was integrated to the IVUS analysis software (QIvus, Medis, Leiden, The Netherlands) which allowed automated characterisation of four tissue types: fibrous tissue (green), fibro-fatty (light green), dense calcium (white) and necrotic core (NC, red).

#### OCT imaging

OCT imaging was performed with a C7XR, an OPTIS ^TM^ (St-Jude Medical, Westford, MA, USA) or a Lunawave Fourier Domain system (Terumo Corp, Tokyo, Japan) depending on the availabilities at the patient recruitment centres. OCT pullback was performed using automated pullback device at a constant speed (range 18-40mm/s) during continuous injection of contrast medium to clear the blood (frame rates: 180fps for the OPTIS, 100fps for the C7XR and 160fps for the Lunawave system). Images were saved for offline analysis that was undertaken using a proprietary software (QCU-CMS, Medis, Leiden, The Netherlands).

In the OCT images, the most proximal and distal side branch located outside the stented segment that was visible in both OCT and in coronary angiography were identified and used to define the segment of interest. OCT segmentation was performed at every 0.4mm intervals (except for the Lunawave system at 0.375mm). The lumen borders of the stented and non-stented segment were detected while, in the stented segment the stent borders were also annotated by connecting the hyperintense signal of the metallic struts. Neointima was defined as the tissue between the lumen and stent border. Neoatherosclerosis was defined as neointimal tissue with lipid (low signal intensity region with diffuse borders and increased signal attenuation) or calcific tissue (heterogenous region with low backscattering and well-defined borders).^14^ In each frame where lipid or calcific tissue is present, the lipid and calcific area were estimated. Macrophages were characterised by signal-rich regions with high signal attenuation behind them.

### Coronary artery reconstruction

The segment of interest in native and stented vessels was reconstructed using two methodologies: the first, the single vessel reconstruction (SVR) method, was able to reconstruct only the main vessel while the second, complete vessel reconstruction (CVR) incorporated side branches with diameter ≥1mm in the final geometry (Figure 1).

**Figure 1.**
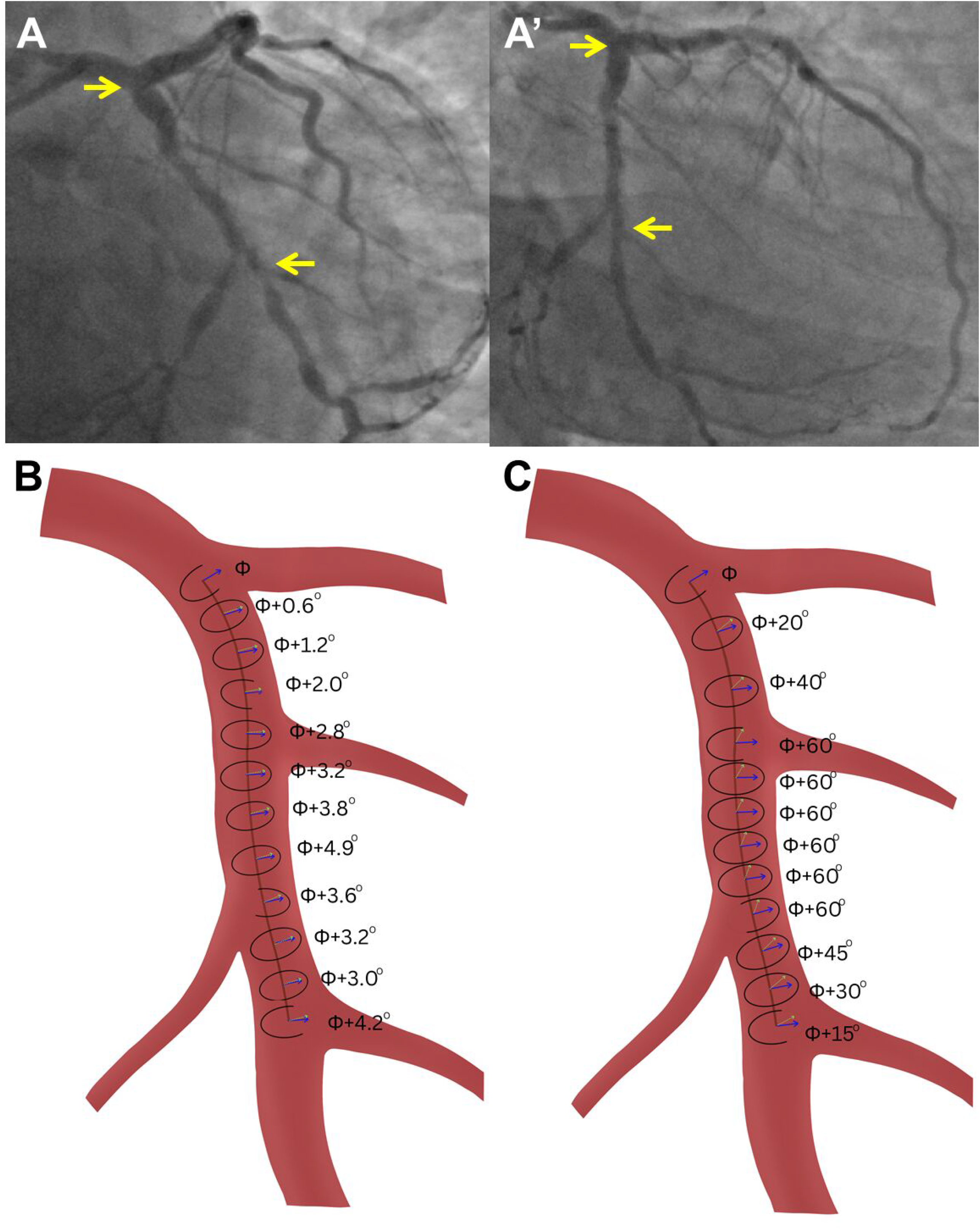
Case example of a vessel reconstruction using the SVR and CVR methodologies. The top panel (A and A’) shows two angiographic projections of a left circumflex artery, the start and end of the reconstructed segment is indicated with yellow arrows. The bottom left panel (B) shows the 3D geometry reconstructed with the SVR methodology. In this method, each IVUS contour is placed perpendicularly onto the vessel centreline at equidistant points and the estimated relative axial twist by the sequential triangulation algorithm is shown. The bottom right panel (C) shows the 3D geometry reconstructed by the CVR methodology. Here, the IVUS contour that portrays the side branches are placed in the corresponding location and rotated to match the origin of the side branches. Linear interpolation is used to estimate the location and orientation of the contours in between the side branches as shown. This approach resulted in larger relative axial twist and variable distances between contours.

#### SVR method

A well-established methodology introduced to retrospectively fuse IVUS or OCT with angiographic data acquired during routine coronary angiography was used to reconstruct the main vessel.^15^ This approach includes: 1) the extraction of the lumen centreline for the segment of interest from two end-diastolic angiographic projections, 2) the placement of the segmented IVUS and OCT borders perpendicularly onto the vessel centreline at equidistant points 3) the computation of the relative axial twist of the intravascular imaging data using the sequential triangulation algorithm and 4) the estimation of the absolute orientation of the first IVUS or OCT frame using a methodology that relies on the comparison of the silhouette of the lumen in the angiographic projections with the back-projected reconstructed model. The angle of the first frame at which the best matching is achieved corresponds to the correct absolute orientation.

The IVUS and X-ray imaging data acquired at baseline and follow-up were used to reconstruct the segment of interest at these two time points while, the OCT data at the time of the event was used to reconstruct two models the lumen geometry at the time of the event and the lumen geometry post stent implantation. The latter reconstruction was based on the assumption that the stent borders in the stented segment and the lumen borders in the non-stented segment at the time of the event corresponded to the lumen immediately after stent implantation.

#### CVR method

The segments of interest were also reconstructed using the methodology introduced by Li et al which allows incorporation of side branches into the 3D coronary artery geometry.^9^ The first step includes QCA of the segment of interest and its side branches with diameter ≥1mm in two end-diastolic angiographic projections (>25° apart) and generation of the corresponding 3D models using the QAngioXA 3D Research Edition 1.0 software (Medis, Leiden, The Netherlands). The 3D QCA models of the main vessel and side branches are then fused to generate the complete geometry of the vessel. The segmented IVUS/OCT images are then fused with the 3D QCA data. More specifically, the frames that portray the side branches are placed in the corresponding location on the 3D model of the main vessel and each one is then rotated so as the origin of the side branches in the IVUS/OCT frame to correspond to the origin of the side branch in the QCA reconstruction. Then linear interpolation is applied to estimate the location and absolute orientation of the in-between frames. Finally, the 3D-QCA model of the main vessel is replaced with the 3D-IVUS/OCT lumen and vessel wall borders that together with the 3D QCA models of the side branches constitute the final geometry for CFD analysis.

### Computational fluid dynamics analysis

To ensure a smooth transition and a fully developed flow when it enters the reconstructed vessel, fixed length extensions of 15mm were added to the inlet and outlet of each model. The lumen and vessel wall models reconstructed with the two approaches were then meshed using ANSYS ICEM 19.0 to generate a tetrahedral volume mesh with five layers of prism elements at the vessel wall. The lumen models reconstructed by the IVUS-VH data acquired at baseline in the IBIS 4 study, and the models that corresponded to the lumen surface at baseline in the stented segments were then imported into ANSYS CFX 19.0 to perform pulsatile flow simulations and calculate the TAWSS and multidirectional WSS by solving the Navier Stokes equations.

Blood was assumed to be non-Newtonian its behaviour was modelled using the Quemada equation; blood density was assumed of 1060kg/m^3^ and its viscosity varied depending on shear rate and haematocrit.^16^ The arterial wall was considered to be rigid, no-slip conditions were applied to the lumen surface while zero pressure conditions were imposed at the outlet. The mean blood flow velocity was estimated as the ratio of: (model length x cine frame rate on angiography) / average number of frames required for the contrast media to pass through the reconstructed segment.^3^ The patient-specific velocity values were combined with generic velocity waveforms reported in the literature to account for the differences between the phasic coronary blood flow in the left and right coronary arteries.^17, 18^ The flow division between the main branch and side branch(es) were calculated according to model proposed by Van der Giessen et al.^19^ The simulation was carried out over three full cardiac cycles. The first two cycles were used for initialization and the third to compute the WSS parameters. Each cardiac cycle was divided into 1000 timesteps.

In each model the TAWSS, which is the WSS averaged over a cardiac cycle, the oscillatory shear index (OSI), introduced to assess flow reversal in pulsatile flow during the cardiac cycle, the relative residence time (RRT) which represent the relative time that a blood particle resides at a distinct location at the arterial wall, the transverse WSS (TransWSS) that reflects the amount of flow perpendicular to the main direction of flow and the cross-flow index (CFI) which is the transWSS normalised by the instantaneous WSS were computed as previously described.^20^

### Data post-processing

The two reconstruction methodologies have differences in the steps that are followed for the generation of the 3D models and in particular, in the position of the borders detected on intravascular images onto the lumen centreline and in the estimation of their rotational orientation. This is likely to affect the 3D model geometry and the longitudinal and circumferential distribution of the plaque. To quantify these differences, we compared the rotational orientation of the intravascular imaging data in the models derived by the two methods. Moreover, we estimated the differences in the location of each intravascular image borders onto the lumen centreline in the models reconstructed by the two methods.

The IVUS-based reconstructions were split in 3mm segments and for each segment we estimated the following variables: the mean lumen, EEM, plaque area and plaque burden (PB, calculated as: 100 × mean plaque area/mean EEM area), and plaque composition (the fibrotic tissue, fibrofatty tissue, NC tissue and calcific tissue areas), derived from IVUS–VH and the percentage of each plaque component. In the OCT-based models analysis focused only in the stent segment; this was split in 1.5mm segments, and for each segment the mean lumen area at follow-up, stent area (that corresponds to the baseline lumen area), neointima area and neointima burden (calculated as: 100 × mean neointima area/mean stent area), and neoatherosclerotic lipid core and calcific area and burden were estimated and the incidence of macrophages were reported.

In addition, for each of the IVUS and OCT segments of the baseline models the minimum predominant TAWSS were estimated and defined as the minimum averaged TAWSS in an arc of 90°.^3, 4^ Moreover, the OSI, RRT, TransWSS and CFI at the 90° arc with the minimum TAWSS were computed. Similarly, to the TAWSS, we estimated the maximum predominant OSI, RRT, TransWSS and CFI defined as the maximum averaged variable in an arc of 90°.

In the IVUS models we have excluded the first and last 3mm segment from the analysis to minimise the effect of the transition from model extension to the realistic geometry and vice versa. Similarly in the OCT models we excluded the first and last 1.5mm segment of the stented segment to minimise the effect of the transition from the lumen to the stent contour.

### Statistical analysis

Continuous variables are presented as median (interquartile range) and categorical as absolute values and percentages. The association between lumen and plaque area and PB and NC burden at follow-up in native segments and neointima area and burden in stented segments and TAWSS and multi- directional WSS were modelled using Bayesian proportional odds model with random effect.^21^ The baseline lumen, plaque area and PB and NC burden were included into the corresponding models. In addition, Bayesian binary logistic regression modelling with random effect was applied to study the association between WSS metrics and disease progression. Univariate analysis was performed for all the above models with the studied vessel been used as random effect. A Markov Chain Monte Carlo method was run with 4 chains and 4000 iterations.

The association between outcomes and TAWSS and multi-directional WSS were quantified by regression coefficient and 95% credible interval (CrI). Regression coefficients were presented over interquartile ranges for predictors based on posterior mode parameters. Non-informative priors were used for regression coefficient. All WSS metrics and other continuous predictors were transformed by using restricted cubic spline (3 knots) to capture the non-linearities.

The variables entered in the univariate analyses were used to build multivariate models to predict the follow-up lumen and plaque area and PB and NC burden in the IVUS reconstructions and neointima area and burden in the OCT reconstructions. The Variance Inflation Factor (VIF) was measured to evaluate multicollinearity among the predictors used in the models; a value greater than 10 was accepted as a serious indication of multicollinearity. In these models the predictors were taken with restricted cubic spline transformation to capture nonlinearities; in this case, multicollinearity can be safely ignored.^22^ A multivariate model was also built to predict disease progression; in this model the OSI and CFI estimations were included in the model as principal component (PC) due to serious multicollinearity (VIF>10), the risk of overfitting due to low number of outcome and the possibility of inflating the degrees of freedom with use of restricted cubic spline.

Multivariate models’ performances were assessed by explained variation (EV, fraction of explained variation in the outcome by model) and C-statistics (or area under the curve, AUC). Finally, a mixed linear regression model was used to compare WSS in segments with and without neoatherosclerotic tissue in stented segment with the studied vessel being used as random intercepts. All statistical analyses were performed by R software v. 4.2.2 (R statistical software, Institute for Statistics and Mathematics, Vienne, Austria) using “rmsb”, “car”, “lme4” and “ggplot2” packages.

## Results

### Intravascular ultrasound models

#### Studied patients

Of the of the 50 patients (n=88 vessels) that were assessed by IVUS imaging at baseline and had biplane angiography, 17 patients (n=19 vessels) fulfilled the inclusion criteria and included in the final analysis. The mean age of studied patients was 52 (47-63) years, and 88.2% were males. None of the patients suffered previous myocardial infarction or underwent PCI. The baseline characteristics of the studied population is described in Supplementary Table 1. The segments assessed by IVUS were in the left anterior descending artery (LAD) in 10 cases (52.6%), in the left circumflex artery (LCx) in 7 (36.8%) and in the right coronary artery (RCA) in 2 (10.5%).

#### Comparison of the reconstructed models

Of the 19 IVUS vessels, follow-up data was available in 18 vessels. The mean length of the reconstructed models was 42.8 (30.6–51.3)mm at baseline and 44.9 (30.4-50.3)mm at follow-up; all models had a single side branch. In the SVR approach, the IVUS contours were placed in equidistant points across the lumen centreline. Conversely, in the CVR method, there was a longitudinal displacement of the IVUS contours so the contours portraying side branches can co-localise with their origin in the 3D-QCA models. This was estimated to be -0.3±1.7mm (range -3 to 6mm) in the baseline and -0.2±1.2mm (range -3.5 to 2mm) in the follow-up models. Moreover, significant differences were reported in the relative rotational orientation of the IVUS contours. In the SVR method, the mean relative axial twist was -0.04±0.55° while in the CVR method the mean torsion was 0.18±1.11°, p<0.001 (Supplementary Figure 1).

#### Comparison in wall shear stress distribution

In total, 213 3mm segments were included in the final analysis. The minimum predominant TAWSS, TransWSS and CFI values were higher in the SVR approach compared to the corresponding estimations in the CVR models (Supplementary Table 2). Conversely, there was no difference between the estimations of the two approaches for the OSI and RRT values. A case example is illustrated in Figure 2.

**Figure 2.**
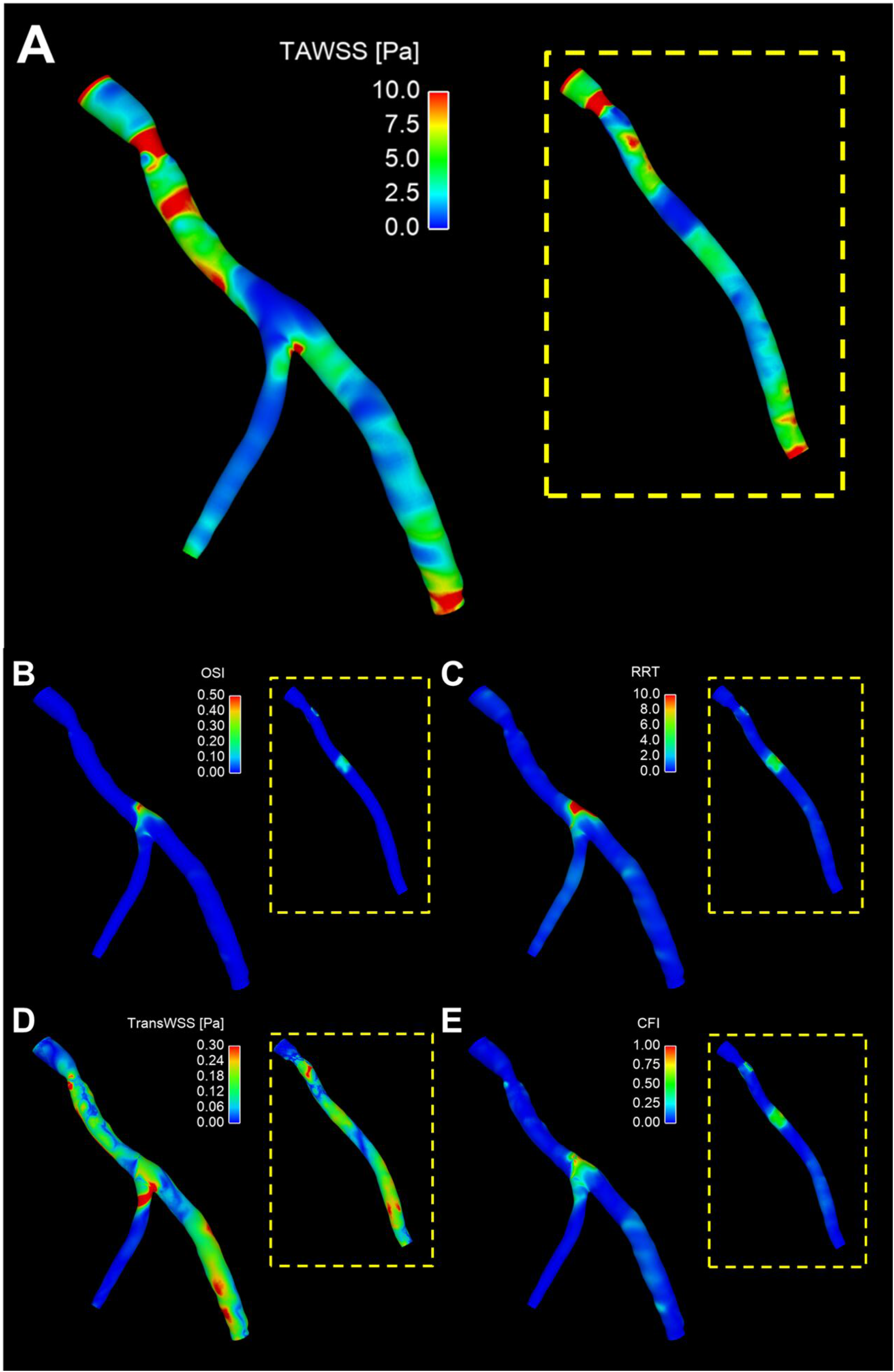
Comparison of wall shear stress distribution. A case example highlighting the differences in the distribution of TAWSS (A), OSI (B), RRT (C), TransWSS (D) and CFI (E) reconstructed using the SVR and CVR methodologies.

#### Association of shear stress with lumen, plaque dimensions and composition at follow-up

The univariate analysis between WSS metrics lumen and plaque area and PB and NC burden is shown in Figure 3. In the SVR model, low TAWSS was associated with small lumen area and increased plaque area at follow-up while RRT was a predictor of increased plaque area, PB and NC burden. High OSI and CFI were associated with increased NC burden at follow-up. In the CVR method, low TAWSS and high multidirectional WSS metrics were associated with smaller lumen area and increased plaque area and PB at follow-up as shown in Figure 3. Moreover, high RRT, TransWSS, and CFI were predictors of a large NC burden at follow-up.

**Figure 3.**
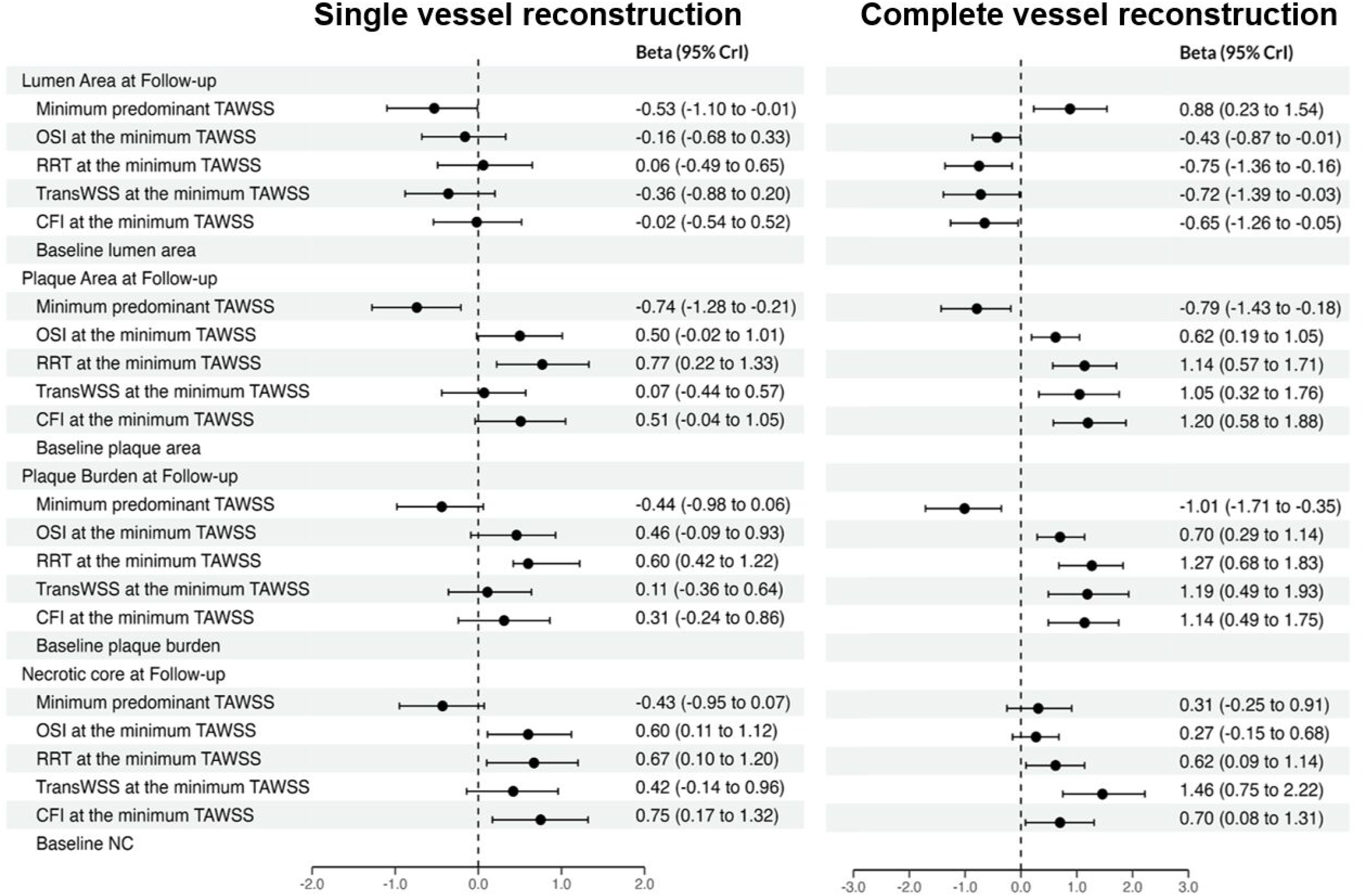
Univariate analysis examining the association of the baseline IVUS estimations and WSS computed by the SVR and CVR methodologies for assessing lumen, plaque dimensions and composition at follow-up.

In the multivariate models, the proportion of EV were significantly lower in the SVR than the CVR method for the follow-up lumen and plaque area and PB but there was no difference between the two methods for the NC burden (Figure 4).

**Figure 4.**
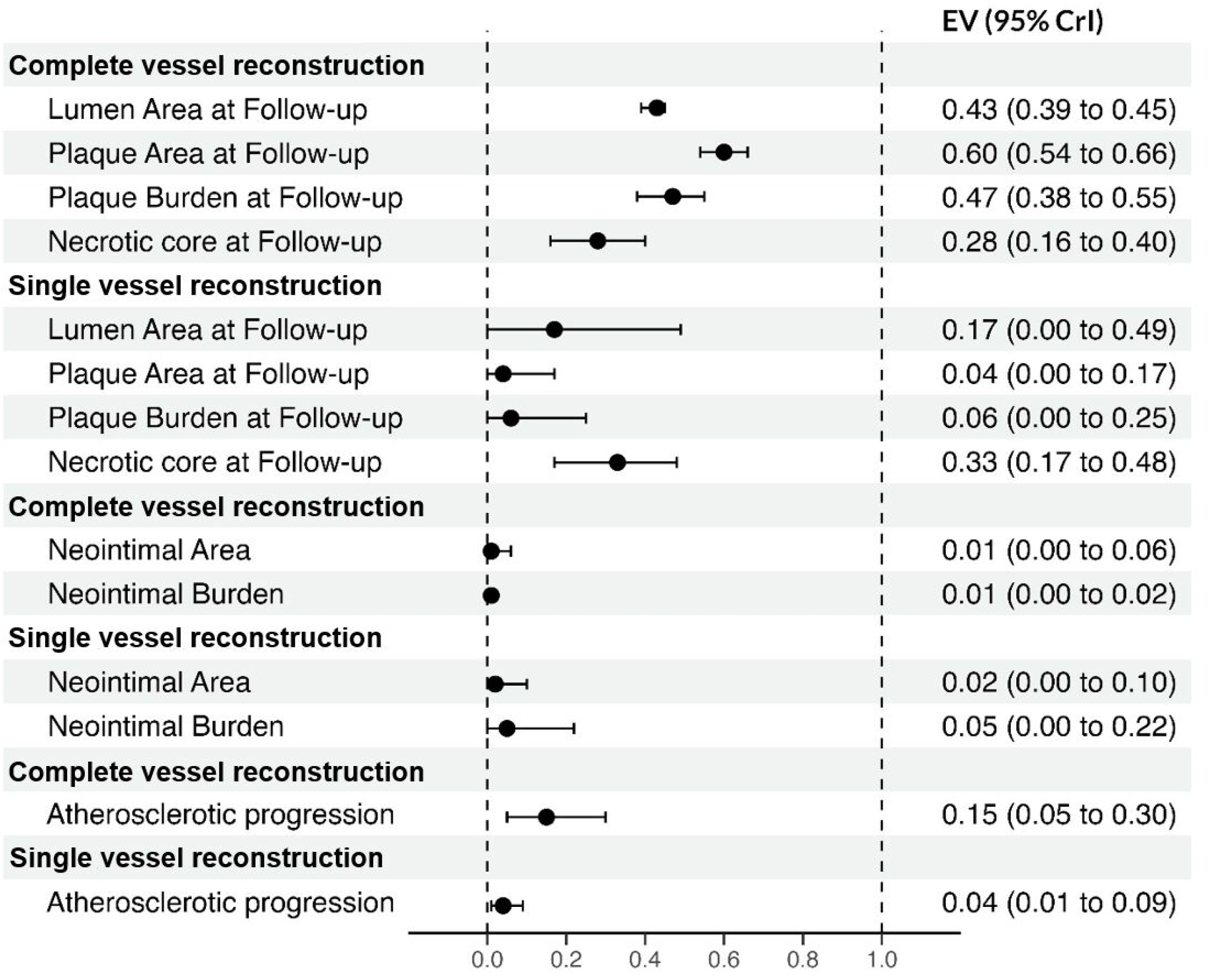
Multivariate models’ performances expressed using the proportion of explained variation (EV) metric for predicting the lumen plaque area PB and necrotic core burden, atherosclerotic disease progression and neointima area and burden. CVR appeared superior compared to SVR with higher proportion of EV for lumen, plaque area and burden and overall atherosclerotic disease progression but no difference was noted between the two methodologies for assessing necrotic core burden at follow-up. In stented segments, both methodologies showed limited efficacy in assessing neointima area and burden.

#### Prediction of disease progression

In the SVR approach, none of the WSS metrics were associated with disease progression at follow-up in univariate analysis. Conversely, in the CVR method, low TAWSS and high OSI, RRT and CFI were predictors of disease progression (Figure 5).

**Figure 5.**
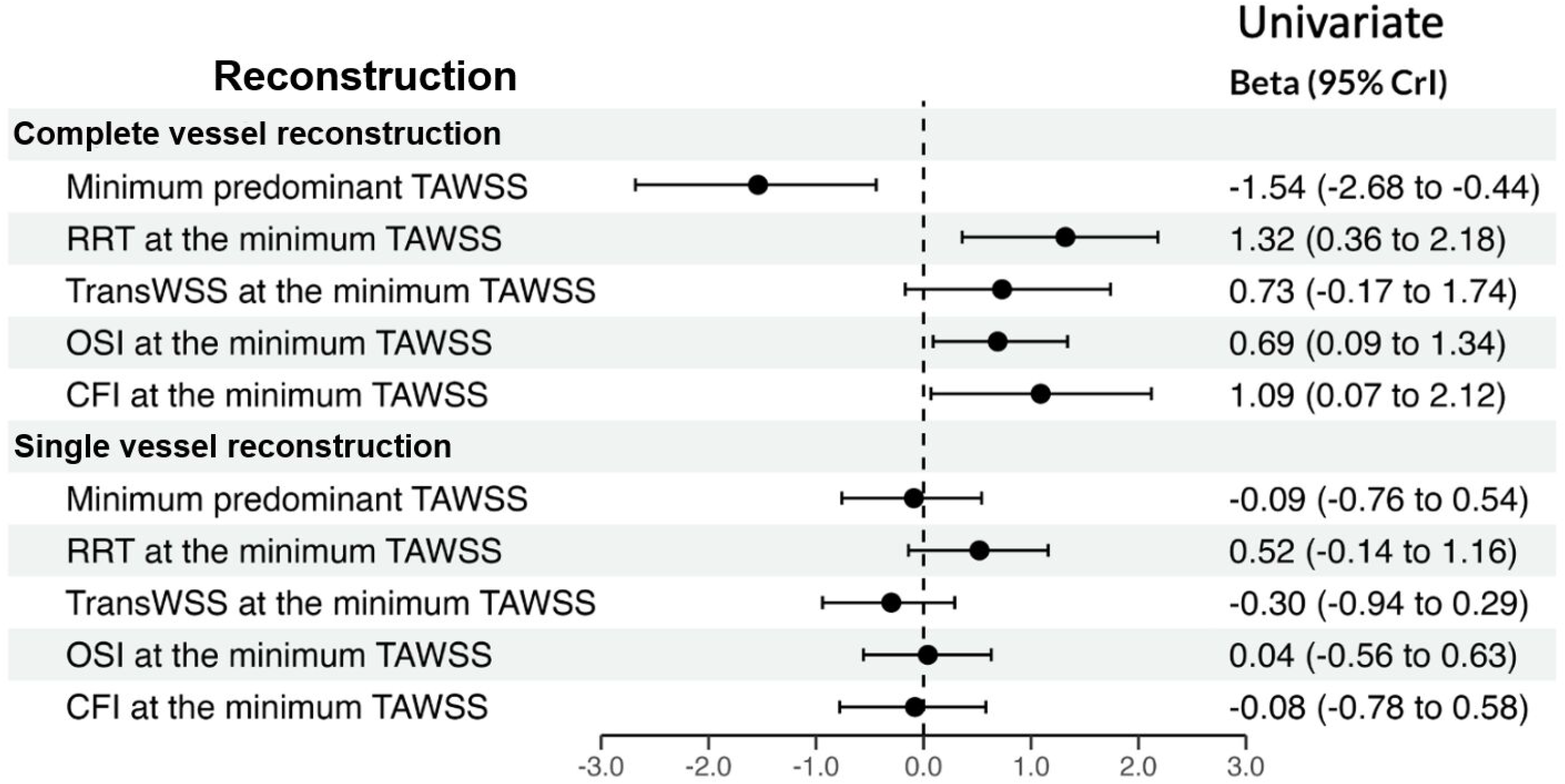
Univariate analysis of atherosclerotic disease progression. In CVR methodology, low TAWSS and high OSI, RRT and CFI were predictors of atherosclerotic disease progression while in SVR none of the WSS metrics were predictors of atherosclerotic disease progression.

The model built by the WSS metrics estimated by the SVR approach had a significantly lower proportion of EV than the model generated by the CVR estimations for atherosclerotic disease progression (Figure 4). The AUC of the model built from the CVR estimations was significantly higher than that of the model including SVR estimations [0.714, (95% CrI 0.679 – 0.729) vs 0.554, (95% CrI 0.528 – 0.578)], as shown in Figure 6.

**Figure 6.**
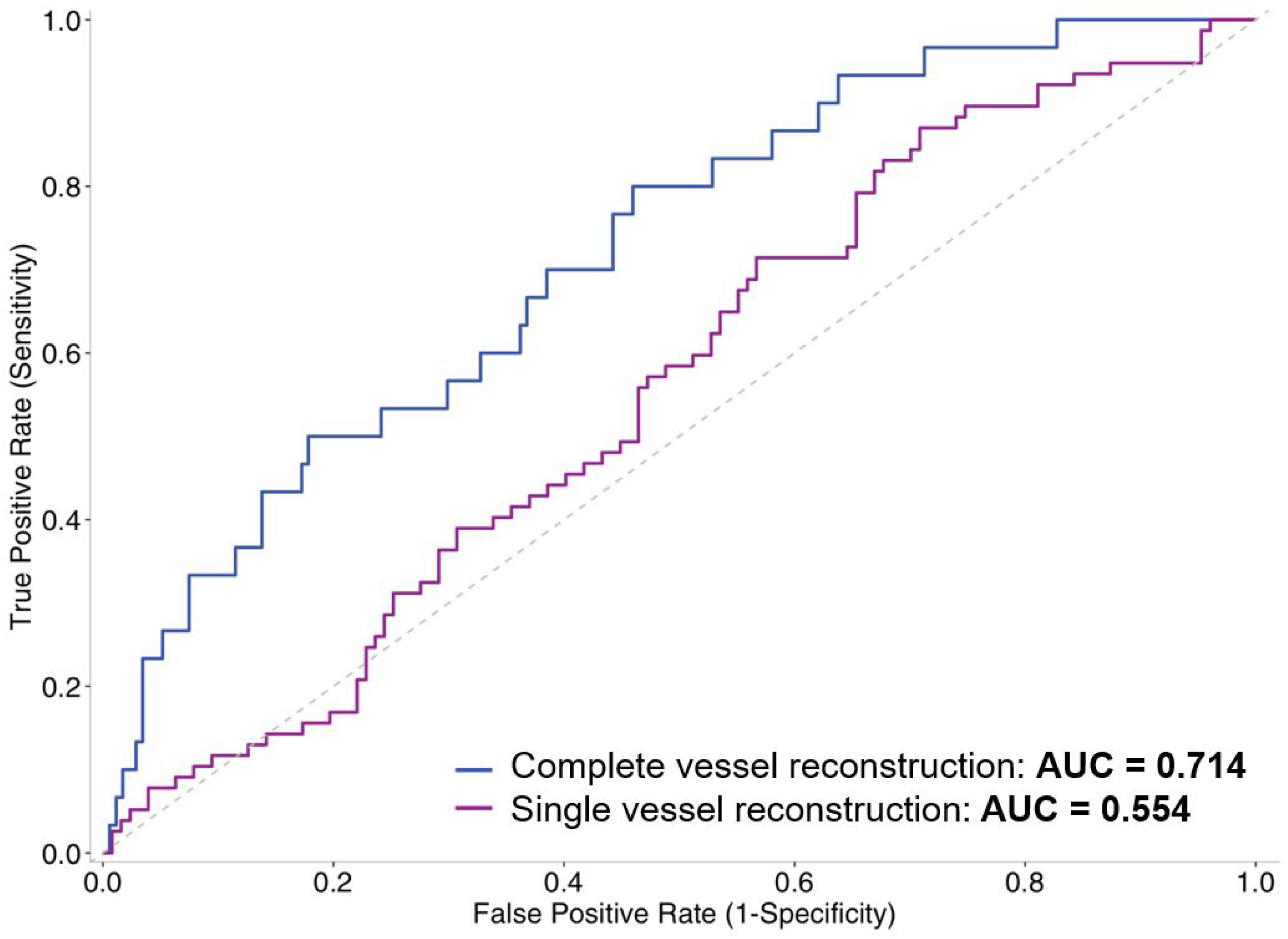
Accuracy of the SVR and CVR methodologies in predicting atherosclerotic disease progression. The model built from the CVR WSS estimations had a higher area under the curve (AUC: 0.714; 95%CI 0.679–0.729) in predicting atherosclerotic disease progression compared to the model built by the SVR WSS (AUC 0.554; 95% CI 0.528–0.578).

### Optical coherence tomography models

#### Studied patients

Thirteen patients that had PCI 15 years ago and presented with an acute coronary syndrome attributed to a neoatherosclerotic lesion, that had at least one side branch with diameter >1mm within the stented segment were included in the analysis; 8 models had a single side branch and the remaining 5 two side-branches. The baseline characteristics of the studied population are presented in Table 1. The mean age of included patients was 61 (57-66) years, 5 (31.5%) patients had hypertension and 10 (76.9%) were diabetic. Most of the studied segments were in the LAD (8, 61.5%), followed by LCx (3, 23.1%) and RCA (2, 15.4%).

#### Comparison of the reconstructed models

The mean length of the reconstructed segments and stents were 34.0 (30.8-38.4)mm and 28.4 (21.7- 35.6)mm, respectively. The longitudinal displacement of the OCT contours in the CVR method ranged from -4 to 8mm (mean 0.7±3.3mm) and was higher than reported in the IVUS-based reconstructions (p=0.028). Similar to the IVUS models, there was a significant difference in the relevant rotation of the OCT contours in the CVR compared to the SVR approach (0.56±2.08° vs 0.00±0.06°, p=0.031).

#### Comparison of shear stress distribution

In total 244 1.5mm segments were included in the final analysis. The TAWSS and multidirectional WSS values are illustrated in Supplementary Table 3. The TAWSS, and transWSS were higher and the OSI, RRT and CFI were lower in the SVR method compared to CVR method.

#### Association of shear stress with neointima area and burden

The association between baseline WSS indices and neointima area and burden are summarised in Figure 7. In the SVR method, all the multidirectional WSS metrics except the TAWSS were associated with neointima area while TransWSS and CFI were related to the neointima burden at follow-up. Conversely in the CVR approach all the WSS metrics including the TAWSS were associated with the neointima area at baseline whereas TransWSS and CFI predicted neointima burden at follow-up.

**Figure 7.**
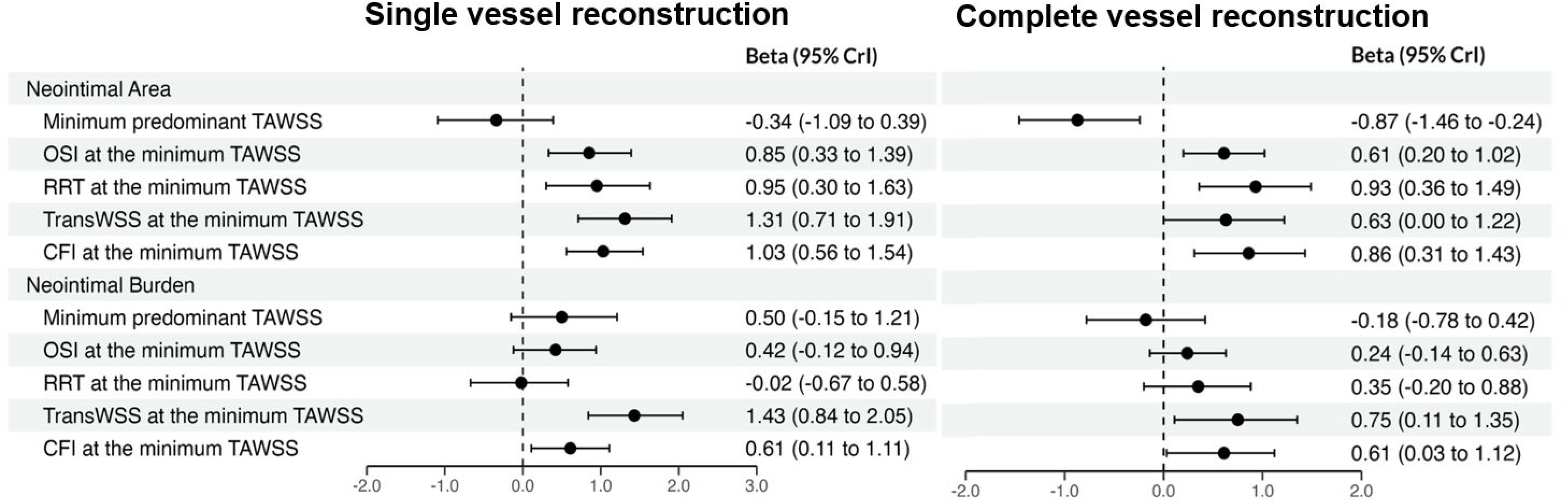
Association of baseline WSS metrics computed by the SVR and CVR methodologies and neointima area and burden at follow-up OCT imaging. All the WSS metrics computed by the CVR and the multidirectional WSS derived by the SVR method were associated with the neointima area at follow-up. Conversely, neointima burden at follow-up was associated with only the baseline high TransWSS and CFI estimated in both CVR and SVR models.

The models built by the WSS variables estimated by the SVR and CVR method had a low proportion of EV for the neointima area and burden indicating that both approaches have a limited efficacy in predicting neointima formation (Figure 4).

#### Shear stress and neoatherosclerotic lesion formation

In the models reconstructed using the SVR method, neoatherosclerosis was identified in 51 (20.5%) segments of which 33 (13.5%) had lipid and 18 (7.4%) calcific tissue while macrophages were seen in 24 segments (9.8%). In the CVR method, the incidence of neoatherosclerosis was 20.9% (n=52), of lipid 11.9% (n=29), of calcific tissue 9.4% (n=23) and of macrophages in 10.2% (n=25). In view of the low incidence of lipid, calcific tissue and of macrophages in the reconstructed geometries, we only compared WSS metrics in segments with and without neoatherosclerosis.

In the models reconstructed using the SVR approach, baseline OSI, RRT and CFI were higher in segments that exhibited neoatherosclerosis at follow-up but there was no difference in TAWSS and TransWSS values between segments with and without neoatherosclerosis. Results were different in the CVR method; the OSI, RRT, TransWSS and CFI values were higher in segments that exhibited neoatherosclerosis at follow-up, while there was no difference in the TAWSS values between segments with and without neoatherosclerosis (Supplementary Figure 2).

Blood flow simulation in models reconstructed using the SVR method required 1 hour per vessel compared to 6 hours per vessel in CVR models (2x16 cores, 128GB RAM, Inter Xeon E5-2640 @2.6GHz, Intel Corporation, Santa Clara, CA, USA).

## Discussion

The present study, for the first time, evaluated the distribution of TAWSS and multidirectional WSS indices in models reconstructed using a SVR method and CVR approach that incorporates vessel side branches in native and stented segments and evaluated the efficacy of the computed WSS metrics in predicting disease progression. We found that 1) there are significant geometrical differences in the reconstructed models; the CVR method requires longitudinal displacement of the IVUS and OCT contours in the lumen centreline and higher degree of rotation between contours for accurate co-registration to the origin of the side branch in intravascular imaging and QCA compared to the SVR, 2) the incorporation of side branches resulted in lower TAWSS and different multidirectional WSS distribution, 3) TAWSS and multidirectional WSS enabled more accurate prediction of disease progression in native vessels in the CVR method and 4) that more WSS metrics derived by the CVR approached were predictors of neointima area and neoatherosclerotic lesion formation than those computed by the SVR approach.

Two previous reports have examined the impact of side branches in flow patterns. Li et al,^9^ who introduced for the first time a method that incorporated the side branches in vessel geometry demonstrated that the inclusion of side branches resulted in a reduction of the mean WSS by 4.64Pa. However, this study included lesions with advanced disease as the mean Pd/Pa across these lesions was 0.85 and therefore it is likely these findings to do not apply to segments with mild stenoses. Similar findings were reported by Giannopoulos et al^23^ who assessed WSS in five hearts imaged *ex vivo* using computed tomography angiography and showed that the exclusion of side branches significantly overestimated WSS by 2.26±3.94Pa and that this methodology reduced the ability of the reconstructed models to predict regions with pathological low WSS. Both studies however performed steady flow simulation and thus they did not assess the effects of side branches on multidirectional WSS and did not examine whether the WSS metrics derived in CVR models allow better prediction of plaque progression and future cardiac events.

The present analysis was designed to address these limitations but also to identify differences in the geometry of models reconstructed by the two approaches. We found that the CVR method requires longitudinal displacement and larger rotation of the intravascular imaging borders. In the IVUS-based reconstructions, the longitudinal displacement was smaller compared to the OCT-based models. This can be explained by the fact that the IVUS images, included in vessel reconstruction, were acquired at the end-diastole – i.e., at the same phase of the cardiac cycle with the X-ray images used for QCA analysis – and thus the longitudinal displacement of the IVUS borders in these models was performed only because of possible foreshortening of a part of the reconstructed segment in X-ray angiography. Conversely, in the OCT-based reconstructions, the included OCT images were obtained during the entire cardiac cycle and therefore it is essential to correct for the relative backward-forward motion of the OCT catheter in the vessel during the cardiac cycle that has been reported up to 5.5mm.^24^

We also found that the rotation of the intravascular image borders is larger in the CVR compared to the estimations of the sequential triangulation algorithm used in the SVR method. However, in silico and experimental studies have shown that the relative axial twist of the intravascular images is limited and is overestimated by the sequential triangulation algorithm.^25–27^ A possible explanation of the larger relative twist noted in CVR method is the fact that often not all the side branch origins are clearly seen in X-ray images resulting in potential errors in the rotational orientation of intravascular images.

The differences in the placement of intravascular images on lumen centreline and in their rotational orientation, as well as the inclusion of the side branches in CVR method resulted in significant differences in WSS distribution in the models reconstructed by the two methodologies. We found that the SVR approach overestimated minimum predominant TAWSS by 0.88Pa in the IVUS-based and by 0.64Pa in the OCT-based reconstructions compared to the CVR method. These differences are smaller to the ones reported in the literature. This should be attributed to the fact that previous reports included diseased segments and also focused on the mean TAWSS values, while this analysis included native segments with minor atheroma (the mean PB at baseline in IVUS-based models was 45.3%) and segments with a large lumen post stent implantation. In addition, we estimated the minimum predominant TAWSS that appear capable to detect plaques prone to progress and cause events.^2–4^

In contrast to previous reports, we performed pulsatile flow simulation and examined for the first time the effects of side branch reconstruction in multidirectional WSS that are instigators of plaque progression.^20, 28^ Coronary bifurcations influence WSS distribution causing flow disturbances at the origin of the daughter branches and an unfavourable haemodynamic milieu that promotes atherosclerotic disease progression.^10^ The flow patterns in these locations cannot be accurately assessed by the models generated by the SVR approach which cannot also take into account flow division in these segments resulting in differences in the multidirectional WSS estimations. In the IVUS-based reconstructions the SVR approach overestimated TransWSS and CFI while in the OCT-based models it overestimated TransWSS and underestimated OSI and RRT.

The limited efficacy of the SVR approach to assess WSS distribution influenced its efficacy in predicting disease progression. The baseline IVUS variables and WSS metrics estimated by the CVR enabled better prediction of these lumen, plaque area and PB at follow-up than the SVR approach as shown by the proportion of EV. Moreover, the efficacy of the WSS metrics in detecting segments that exhibited disease progression at follow-up was significantly higher in CVR than the WSS computed by the SVR approach.

Similarly, to the native segments in the stented segments more WSS metrics derived by the CVR approach were associated with neointima area and burden than the estimations of the SVR. However, in this dataset both CVR and SVR had a limited efficacy in predicting vessel wall healing at follow-up. This should be attributed to the fact that we assumed in these models that the stent borders at the time of the event corresponded to the lumen borders post stent implantation – as the baseline OCT data was not available – and thus it was not possible to examine in detail the implications of the local haemodynamic micro-milieu, that is determined by strut apposition, on neointimal proliferation, and to the large time interval between stent implantation and OCT imaging in this group (7.4±4.8 years). *Limitations*

Several limitations of the present analysis should be acknowledged. Firstly, the number of included native segments is relatively small; therefore, it was not possible to examine the value of WSS metrics in detecting disease progression at lesion-level and vulnerable plaques. This should be explored in future large scale prospective studies. Similarly, the 13 stented segments of the present analysis did not allow assessment of the effects of WSS on lipid-rich neoatherosclerotic lesion formation and macrophages accumulations. Secondly, as stated above in the stented segments, the OCT data post-stent implantation were not available. To overcome this limitation, it was assumed that the stent borders at the time of the event corresponded to the lumen borders post device deployment. This approach has been extensively used in the past to evaluate the effect of flow patterns on vessel wall healing.^13, 29, 30^ Similarly to previous studies we found an association between WSS and neointima area and burden at the time of the event, but the overall efficacy of the model including the WSS variables to predict neointima proliferation was limited. Therefore, future studies with serial OCT imaging should be conducted to fully examine the value of baseline WSS distribution in predicting vessel healing. Finally, flow in the studied segments was estimated by the TIMI frame count and not by a Doppler wire; this can affect the estimated WSS metrics especially in the models incorporating the side branches where flow division was approximated in our study. Despite these approximations however the CVR method provided WSS estimations that enabled more accurate prediction of disease progression and was strongly associated with neointima characteristics in stented segments

## Conclusion

CVR and incorporation of side branches in 3D vessel geometry is feasible and results in different WSS distribution compared to the SVR approach. This method enables more accurate identification of segments prone to progress in native vessels and provides WSS estimations that are strongly related to neointima proliferation in stented segments. Therefore, his approach should be preferred for evaluating the implications of the local haemodynamic forces in vessel wall pathobiology in native and stented segments despite the fact that it is consuming, computationally challenging and expensive.

**Supplementary Figure 1**. Relative rotational orientation of IVUS contours in a segment located in the LCx reconstructed using the SVR and CVR method. It is apparent that the relative axial twist of the intravascular images is considerably smaller in the SVR (blue) methodology than the mean torsion noted in the CVR (orange) methodology.

**Supplementary Figure 2.** Comparison of baseline WSS metrics computed by the SVR and CVR methodologies in segments with and without neoatherosclerotic tissue at follow-up.

## References

1. Koskinas KC, Chatzizisis YS, Baker AB, Edelman ER, Stone PH and Feldman CL. The role of low endothelial shear stress in the conversion of atherosclerotic lesions from stable to unstable plaque. Curr Opin Cardiol. 2009;24:580–90.

2. Stone PH, Maehara A, Coskun AU, Maynard CC, Zaromytidou M, Siasos G, Andreou I, Fotiadis D, Stefanou K, Papafaklis M, Michalis L, Lansky AJ, Mintz GS, Serruys PW, Feldman CL and Stone GW. Role of Low Endothelial Shear Stress and Plaque Characteristics in the Prediction of Nonculprit Major Adverse Cardiac Events: The PROSPECT Study. JACC Cardiovascular imaging. 2018;11:462–471.

3. Stone PH, Saito S, Takahashi S, Makita Y, Nakamura S, Kawasaki T, Takahashi A, Katsuki T, Nakamura S, Namiki A, Hirohata A, Matsumura T, Yamazaki S, Yokoi H, Tanaka S, Otsuji S, Yoshimachi F, Honye J, Harwood D, Reitman M, Coskun AU, Papafaklis MI and Feldman CL. Prediction of progression of coronary artery disease and clinical outcomes using vascular profiling of endothelial shear stress and arterial plaque characteristics: the PREDICTION Study. Circulation. 2012;126:172–81.

4. Bourantas CV, Räber L, Sakellarios A, Ueki Y, Zanchin T, Koskinas KC, Yamaji K, Taniwaki M, Heg D, Radu MD, Papafaklis MI, Kalatzis F, Naka KK, Fotiadis DI, Mathur A, Serruys PW, Michalis LK, Garcia-Garcia HM, Karagiannis A and Windecker S. Utility of Multimodality Intravascular Imaging and the Local Hemodynamic Forces to Predict Atherosclerotic Disease Progression. JACC: Cardiovascular Imaging. 2020;13:1021–1032.

5. Kilic Y, Safi H, Bajaj R, Serruys PW, Kitslaar P, Ramasamy A, Tufaro V, Onuma Y, Mathur A, Torii R, Baumbach A and Bourantas CV. The Evolution of Data Fusion Methodologies Developed to Reconstruct Coronary Artery Geometry From Intravascular Imaging and Coronary Angiography Data: A Comprehensive Review. Frontiers in cardiovascular medicine. 2020;7:33.

6. Bourantas CV, Ramasamy A, Karagiannis A, Sakellarios A, Zanchin T, Yamaji K, Ueki Y, Shen X, Fotiadis DI, Michalis LK, Mathur A, Serruys PW, Garcia-Garcia HM, Koskinas K, Torii R, Windecker S and Räber L. Angiographic derived endothelial shear stress: a new predictor of atherosclerotic disease progression. European heart journal cardiovascular Imaging. 2019;20:314–322.

7. Toutouzas K, Chatzizisis YS, Riga M, Giannopoulos A, Antoniadis AP, Tu S, Fujino Y, Mitsouras D, Doulaverakis C, Tsampoulatidis I, Koutkias VG, Bouki K, Li Y, Chouvarda I, Cheimariotis G, Maglaveras N, Kompatsiaris I, Nakamura S, Reiber JH, Rybicki F, Karvounis H, Stefanadis C, Tousoulis D and Giannoglou GD. Accurate and reproducible reconstruction of coronary arteries and endothelial shear stress calculation using 3D OCT: comparative study to 3D IVUS and 3D QCA. Atherosclerosis. 2015;240:510–9.

8. Murata N, Hiro T, Takayama T, Migita S, Morikawa T, Tamaki T, Mineki T, Kojima K, Akutsu N, Sudo M, Kitano D, Fukamachi D, Hirayama A and Okumura Y. High shear stress on the coronary arterial wall is related to computed tomography-derived high-risk plaque: a three-dimensional computed tomography and color-coded tissue-characterizing intravascular ultrasonography study. Heart Vessels. 2019;34:1429–1439.

9. Li Y, Gutiérrez-Chico JL, Holm NR, Yang W, Hebsgaard L, Christiansen EH, Mæng M, Lassen JF, Yan F, Reiber JH and Tu S. Impact of Side Branch Modeling on Computation of Endothelial Shear Stress in Coronary Artery Disease: Coronary Tree Reconstruction by Fusion of 3D Angiography and OCT. Journal of the American College of Cardiology. 2015;66:125–35.

10. Gijsen F, Katagiri Y, Barlis P, Bourantas C, Collet C, Coskun U, Daemen J, Dijkstra J, Edelman E, Evans P, van der Heiden K, Hose R, Koo BK, Krams R, Marsden A, Migliavacca F, Onuma Y, Ooi A, Poon E, Samady H, Stone P, Takahashi K, Tang D, Thondapu V, Tenekecioglu E, Timmins L, Torii R, Wentzel J and Serruys P. Expert recommendations on the assessment of wall shear stress in human coronary arteries: existing methodologies, technical considerations, and clinical applications. European heart journal. 2019;40:3421–3433.

11. Stone PH, Coskun AU and Prati F. Ongoing Methodological Approaches to Improve the In Vivo Assessment of Local Coronary Blood Flow and Endothelial Shear Stress: The Devil Is in the Details∗. Journal of the American College of Cardiology. 2015;66:136–138.

12. Räber L, Taniwaki M, Zaugg S, Kelbæk H, Roffi M, Holmvang L, Noble S, Pedrazzini G, Moschovitis A, Lüscher TF, Matter CM, Serruys PW, Jüni P, Garcia-Garcia HM and Windecker S. Effect of high-intensity statin therapy on atherosclerosis in non-infarct-related coronary arteries (IBIS-4): a serial intravascular ultrasonography study. European heart journal. 2015;36:490–500.

13. Torii R, Stettler R, Räber L, Zhang YJ, Karanasos A, Dijkstra J, Patel K, Crake T, Hamshere S, Garcia-Garcia HM, Tenekecioglu E, Ozkor M, Baumbach A, Windecker S, Serruys PW, Regar E, Mathur A and Bourantas CV. Implications of the local hemodynamic forces on the formation and destabilization of neoatherosclerotic lesions. International journal of cardiology. 2018;272:7–12.

14. Stettler R, Dijkstra J, Räber L, Torii R, Zhang YJ, Karanasos A, Lui S, Crake T, Hamshere S, Garcia-Garcia HM, Tenekecioglu E, Ozkor M, Windecker S, Serruys PW, Regar E, Mathur A and Bourantas CV. Neointima and neoatherosclerotic characteristics in bare metal and first- and second-generation drug-eluting stents in patients admitted with cardiovascular events attributed to stent failure: an optical coherence tomography study. EuroIntervention : journal of EuroPCR in collaboration with the Working Group on Interventional Cardiology of the European Society of Cardiology. 2018;13:e1831–e1840.

15. Bourantas CV, Papafaklis MI, Athanasiou L, Kalatzis FG, Naka KK, Siogkas PK, Takahashi S, Saito S, Fotiadis DI, Feldman CL, Stone PH and Michalis LK. A new methodology for accurate 3-dimensional coronary artery reconstruction using routine intravascular ultrasound and angiographic data: implications for widespread assessment of endothelial shear stress in humans. EuroIntervention : journal of EuroPCR in collaboration with the Working Group on Interventional Cardiology of the European Society of Cardiology. 2013;9:582–93.

16. Quemada D. Rheology of concentrated disperse systems II. A model for non-newtonian shear viscosity in steady flows. Rheologica Acta. 1978;17:632–642.

17. Davies JE, Whinnett ZI, Francis DP, Manisty CH, Aguado-Sierra J, Willson K, Foale RA, Malik IS, Hughes AD, Parker KH and Mayet J. Evidence of a dominant backward-propagating “suction” wave responsible for diastolic coronary filling in humans, attenuated in left ventricular hypertrophy. Circulation. 2006;113:1768–78.

18. Hadjiloizou N, Davies JE, Malik IS, Aguado-Sierra J, Willson K, Foale RA, Parker KH, Hughes AD, Francis DP and Mayet J. Differences in cardiac microcirculatory wave patterns between the proximal left mainstem and proximal right coronary artery. Am J Physiol Heart Circ Physiol. 2008;295:H1198–h1205.

19. van der Giessen AG, Groen HC, Doriot PA, de Feyter PJ, van der Steen AF, van de Vosse FN, Wentzel JJ and Gijsen FJ. The influence of boundary conditions on wall shear stress distribution in patients specific coronary trees. Journal of biomechanics. 2011;44:1089–95.

20. Hoogendoorn A, Kok AM, Hartman EMJ, de Nisco G, Casadonte L, Chiastra C, Coenen A, Korteland SA, Van der Heiden K, Gijsen FJH, Duncker DJ, van der Steen AFW and Wentzel JJ. Multidirectional wall shear stress promotes advanced coronary plaque development: comparing five shear stress metrics. Cardiovascular research. 2020;116:1136–1146.

21. Liu Q, Shepherd BE, Li C and Harrell FE, Jr. Modeling continuous response variables using ordinal regression. Stat Med. 2017;36:4316–4335.

22. Harrell FE, Jr. Regression modeling strategies: Springer International Publishing; 2016.

23. Giannopoulos AA, Chatzizisis YS, Maurovich-Horvat P, Antoniadis AP, Hoffmann U, Steigner ML, Rybicki FJ and Mitsouras D. Quantifying the effect of side branches in endothelial shear stress estimates. Atherosclerosis. 2016;251:213–218.

24. Arbab-Zadeh A, DeMaria AN, Penny WF, Russo RJ, Kimura BJ and Bhargava V. Axial movement of the intravascular ultrasound probe during the cardiac cycle: implications for three-dimensional reconstruction and measurements of coronary dimensions. American heart journal. 1999;138:865–72.

25. Wahle A, Prause PM, DeJong SC and Sonka M. Geometrically correct 3-D reconstruction of intravascular ultrasound images by fusion with biplane angiography-- methods and validation. IEEE Trans Med Imaging. 1999;18:686–99.

26. Prause GP, DeJong SC, McKay CR and Sonka M. Towards a geometrically correct 3- D reconstruction of tortuous coronary arteries based on biplane angiography and intravascular ultrasound. International journal of cardiac imaging. 1997;13:451–62.

27. Bourantas CV, Kourtis IC, Plissiti ME, Fotiadis DI, Katsouras CS, Papafaklis MI and Michalis LK. A method for 3D reconstruction of coronary arteries using biplane angiography and intravascular ultrasound images. Comput Med Imaging Graph. 2005;29:597–606.

28. Kok AM, Molony DS, Timmins LH, Ko YA, Boersma E, Eshtehardi P, Wentzel JJ and Samady H. The influence of multidirectional shear stress on plaque progression and composition changes in human coronary arteries. EuroIntervention : journal of EuroPCR in collaboration with the Working Group on Interventional Cardiology of the European Society of Cardiology. 2019;15:692–699.

29. Bourantas CV, Papafaklis MI, Naka KK, Tsakanikas VD, Lysitsas DN, Alamgir FM, Fotiadis DI and Michalis LK. Fusion of optical coherence tomography and coronary angiography - in vivo assessment of shear stress in plaque rupture. International journal of cardiology. 2012;155:e24–6.

30. Papafaklis MI, Bourantas CV, Theodorakis PE, Katsouras CS, Naka KK, Fotiadis DI and Michalis LK. The effect of shear stress on neointimal response following sirolimus- and paclitaxel-eluting stent implantation compared with bare-metal stents in humans. JACC Cardiovascular interventions. 2010;3:1181–9.

31. Bourantas CV, Papafaklis MI, Kotsia A, Farooq V, Muramatsu T, Gomez-Lara J, Zhang YJ, Iqbal J, Kalatzis FG, Naka KK, Fotiadis DI, Dorange C, Wang J, Rapoza R, Garcia-Garcia HM, Onuma Y, Michalis LK and Serruys PW. Effect of the endothelial shear stress patterns on neointimal proliferation following drug-eluting bioresorbable vascular scaffold implantation: an optical coherence tomography study. JACC Cardiovasc Interv. 2014;7:315–24.

32. Radu MD, Raber L, Heo J, Gogas BD, Jorgensen E, Kelbaek H, Muramatsu T, Farooq V, Helqvist S, Garcia-Garcia HM, Windecker S, Saunamaki K and Serruys PW. Natural history of optical coherence tomography-detected non-flow-limiting edge dissections following drug-eluting stent implantation. EuroIntervention. 2014;9:1085–94.

33. Gutierrez-Chico JL, Wykrzykowska J, Nuesch E, van Geuns RJ, Koch KT, Koolen JJ, di Mario C, Windecker S, van Es GA, Gobbens P, Juni P, Regar E and Serruys PW. Vascular tissue reaction to acute malapposition in human coronary arteries: sequential assessment with optical coherence tomography. Circ Cardiovasc Interv. 2012;5:20–9, S1-8.

